# Multi-ancestry tandem repeat association study of hair colour using exome-wide sequencing

**DOI:** 10.1101/2024.02.24.581865

**Authors:** Victoria Fazzari, Ashley Moo-Choy, Mary Anne Panoyan, Cristina L Abbatangelo, Renato Polimanti, Nicole MM Novroski, Frank R Wendt

## Abstract

Hair colour variation is influenced by hundreds of positions across the human genome but this genetic contribution has only been narrowly explored. Genome-wide association studies identified single nucleotide polymorphisms (SNPs) influencing hair colour but the biology underlying these associations is challenging to interpret. We report 16 tandem repeats (TRs) with effects on different models of hair colour plus two TRs associated with hair colour in diverse ancestry groups. Several of these TRs expand or contract amino acid coding regions of their localized protein such that structure, and by extension function, may be altered. We also demonstrate that independent of SNP variation, these TRs can be used to great an additive polygenic score that predicts darker hair colour. This work adds to the growing body of evidence regarding TR influence on human traits with relatively large and independent effects relative to surrounding SNP variation.

## Introduction

Hair colour is a highly polygenic trait, exhibiting considerable variation within and between human populations (1). The pigmentation of human hair is determined primarily by the type and amount of melanin located in the hair follicle (1,2). Melanin is produced in epidermal melanocytes, specifically within organelles known as melanosomes, and subsequently transported to keratinocytes where they play a role in defending against ultraviolet radiation (1). Melanosomes are responsible for synthesizing two forms of melanin: a red or yellow pigment called pheomelanin and a brown or black pigment called eumelanin (1,3). The amount and ratio of eumelanin and pheomelanin largely contribute to hair colour variation amongst individuals (4). Eumelanin content is highest in individuals with black and dark brown hair, and lower in individuals with light brown and blonde hair (1). Pheomelanin is unique as it is only contained in substantial quantities in individuals with red hair (1).

In contrast to the clinal global distribution demonstrated by skin pigmentation, hair colour does not follow such a regular patten. In fact, there is very little variation in hair colour within populations outside of Europe, and even less variation in eye colour within populations outside of Europe, North Africa, the Middle East, Central Asia and South Asia (5). Scalp hair demonstrates low levels of phenotypic variation in populations outside of Europe, with dark brown being the most common pigment (2). There are some deviations from this pattern, for example, in Northern Island Melanesia there is a relatively high prevalence of blonde hair, which has been attributed to one specific allele, the 93C allele of the *TYRP1* gene. The dispersal of the allele throughout the region was traced back to the human colonization of the South West Pacific (6). In Europe, a different pathway is responsible for the occurrence of blonde hair (evidence of convergent evolution); here, blond hair is the result of variation in a regulatory enhancer of the *KITLG* gene, and the rare red hair phenotype is produced by a very specific range of variants of the *MC1R* locus (7).

Recent large-scale genome-wide association studies (GWASs) have highlighted the high polygenicity and heritability of hair colour (3,8). A study involving approximately 300,000 European participants from 23andMe, Inc. and UK Biobank discovered that 123 autosomal and one X-chromosome loci were significantly associated with hair colour (8). The single nucleotide polymorphisms (SNPs) identified within these regions cumulatively explained 26.1%, 24.8%, and 34.6% of black, blonde, and red hair variability, respectively (8). A more recent GWAS performed in the UK Biobank identified 128 putative causal SNPs associated with red hair alone, accounting for approximately 35% of phenotypic variation (9). A separate GWAS study utilizing nearly 350,000 UK Biobank British participants of European descent identified nine genetic variants associated with red hair that accounted for approximately 90% of SNP heritability, with the *a priori MC1R* variants explaining 73% of that sum (3). Moreover, over 200 variants were associated with blond hair and accounted for 73% of the SNP heritability, whereas the loci associated with brown hair accounted for 47% of the SNP heritability (3). Notably, however, studies assessing genetic variation in hair colour have been performed in relatively homogenous populations, with the majority of participants of European descent (3,10,11).

Although studies have demonstrated the functional role and underlying biology of pigmentation pathways for individual genes, such as a loss of function on the MC1R gene is associated with fair, UV-sensitive, and melanoma-prone phenotypes, likely due to defective epidermal melanization and sub-optimal DNA repair (12). The SNPs discovered from recent GWAS of hair colour are largely intergenic (i.e., found between genes rather than within a gene), making it difficult to understand how the polygenic signal for hair colour contributes to pigment-associated biological pathways and mechanisms. This presents several limitations as associated SNPs may not be causal, demonstrate little heritability and do not account for additional genetic architecture that affects hair colour (13). Therefore, by studying other genetic variants, such as tandem repeats (TRs) in exonic regions, additional and potentially more resolved information can be discovered.

TRs are highly polymorphic regions of the genome that comprise a DNA sequence unit that is repeated a variable number of times (14). When TRs appear in a protein-coding region or exon, the number of repetitive elements may modify subsequent translation processes, protein folding and function, and ultimately influence human health and disease. For this reason, TRs are valuable targets for association studies since expansions and contractions of this region can be associated with traits. From a pathogenic perspective, TR expansions relative to contractions are typically associated with monogenic disorders (15). Among complex traits, TR variation is associated with autism spectrum disorder (ASD) (15), glaucoma (16), cancer (16), brain volume (17), human height (18), and blood and serum traits (19). Additionally, our group has previously also demonstrated an effect of the dinucleotide repeat *NCOA6*-[GT]_N_ on multiple pigmentation-related traits in the UK Biobank (20). Despite our prior work on *de novo* coding TRs, no study to date has investigated the entire coding region for TR effects on hair colour.

In this study, whole exome sequences obtained from 141,777 unrelated participants from six ancestry groups (i.e., African, AFR N=2,607; Admixed-American, AMR N=422; Central-South Asian, CSA N=3,765; East Asian, EAS N=1,153; European, EUR N=133,190; and Middle Eastern, MID N=640) were used to identify TR genotypes at 15,947 TRs. These loci were associated with three models of hair colour variation. We identified 16 TRs associated with hair colour, some of which (i) localize to known hair colour regions but have larger effects than hair colour informative SNPs, (ii) fine-map relative to surrounding biallelic variation, (iii) influence 3-dimensional protein structure, and (iv) additively predict hair darkening in a novel approach to constructing a TR PGS. This work adds to the growing body of evidence that TRs influence a wide range of human traits with relatively large and independent effects relative to surrounding SNP variation.

## Results

### TRs associated with hair colour

The UK Biobank has hair colour data for six global population groups (Fig. S1). The largest sample size is for the EUR individuals who will serve as our discovery sample. In the discovery set of 133,163 unrelated EUR participants (90% of the total), we identified 16 TRs with a significant association with hair colour (FDR q<0.05): 12 in the blonde hair analysis, 8 in the red hair analysis, and 11 in the low eumelanin analysis (Figure 1A and Table S1). Six of these loci had significant effects across all three ordinal models: (i) blonde, light brown, dark brown, and black with red hair excluded (3), (ii) red, light brown, dark brown, and black with blonde hair excluded (3), (iii) “low eumelanin” (i.e., red or blonde) hair colour compared to light brown, dark brown, and black hair (Fig. 1B). Of the loci discovered, 66% (8/12) of the blonde hair TRs, 75% (6/8) of the red hair TRs, and 72.7 (8/11) of the low eumelanin TRs replicated in the remaining 10% of EUR participants (Fig. S2 and Table S1). In all models, the most significantly associated of TR was *FAM157C*-[GCA]_N_ (blonde hair lJ=-0.011, P=7.14×10^-88^, red hair lJ=-0.013, P=2.05×10^-217^, low eumelanin lJ=-0.015, P=9.56×10^-153^). There was evidence that this locus exhibits quantitative heterogeneity across models with a significantly lower effect on overall low eumelanin hair than exclusively red hair (heterogeneity P=5.09×10^-5^) or blonde hair (heterogeneity P=6.22×10^-7^). Hair colour genes were enriched for cellular compartments related to melanin storage: melanosome membrane and pigment granule membrane (Fig. 1C and Tables S2-S4). We also detected concordant enrichment of glycosylphosphatidylinositol-anchored proteins across all three hair colour association models (GO:00034235 and GO:0016255; Fig. 1C and Table S2-S4).

**Fig. 1.**
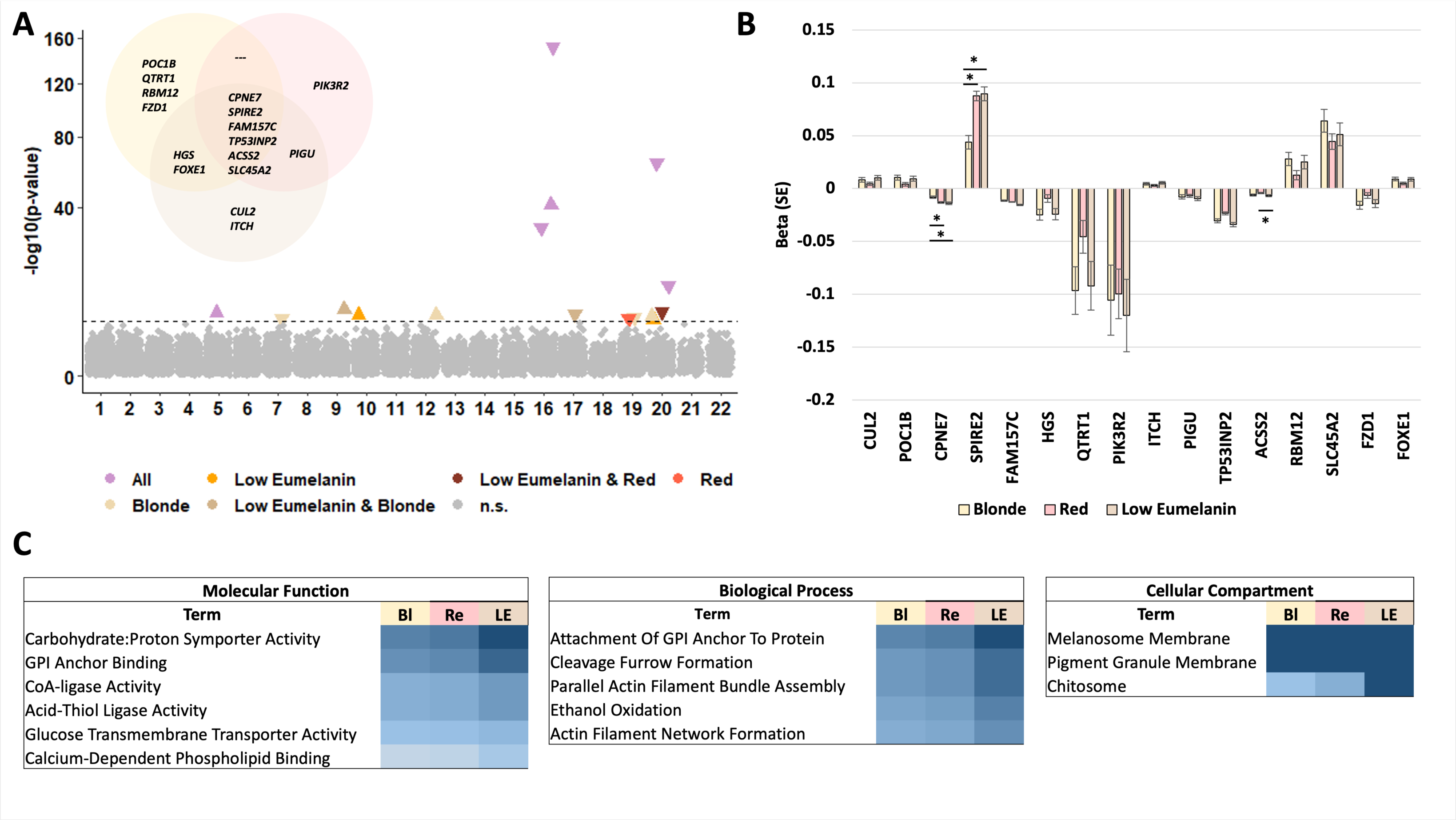
Hair colour TRs in European ancestry participants. Summary of tandem repeats associated with hair colour based on three models: (i) blonde versus light brown, dark brown, and black, (ii) red versus light brown, dark brown, and black, and (iii) low eumelanin (red or blonde) versus light brown, dark brown, and black. Panel A depicts all associated loci in the three models with data point colour representing the models in which the TR met statistical significance (false discovery rate < 5%). Data point shape indicates the effect direction of the TR on hair colour with a upward triangle indicating the TR associates with darker hair colour. Panel B shows the European ancestry effect size of each discovered TR in all three models. Asterisks denote significant differences in effect size after multiple testing correction (P<0.001) determined using two-sided Z-tests. Panel C shows gene set enrichment results for each hair colour model (Blonde=“Bl”, Red=“Re”, low eumelanin=“LE”); all FDR adjusted P<0.05.

### Hair colour TRs in the MC1R region

*FAM157C*-[GCA]_N_ is part of a gene-dense region that includes two additional significantly associated TRs: *CPNE7*-[A]_N_ (blonde hair lJ=-0.008, P=3.04×10^-12^, red hair lJ=-0.013, P=7.39×10^-59^, low eumelanin lJ=-0.014, P=1.65×10^-31^) and *SPIRE2*-[GAG]_N_ (blonde hair lJ=0.044, P=6.39×10^-12^, red hair lJ=0.088, P=6.79×10^-85^, low eumelanin lJ=0.089, P=9.73×10^-42^). Notably, *SPIRE2*-[GAG]_N_ encodes a stretch of glutamate residues in the kinase non-catalytic C-lobe domain of protein spire homolog 2. This region of chromosome 16 is home to the large *MC1R* region with multiple independent SNPs known to influence hair colour (21). Using the program FINEMAP (version 1.4.2) (22), we performed long-range fine-mapping of a 2.6Mb region containing *MC1R* and the three TR-containing genes *FAM157C*, *SPIRE2*, and *CPNE7* (Fig. 2A). LD matrices and variant effect sizes were estimated using the same set of UKB individuals used for TR association testing. For each locus, the maximum number of causal variants was set to 20 (i.e., a maximum of 20 credible sets). In all three hair colour models, we identified 14 variants in the credible set (log_10_(BF) ranged from 3303.25 to 8994.18). Among those were the well-known *MC1R* variants rs1805005, rs1805007, and rs1805008. However, *SPIRE2*-[GAG]_N_ and *CPNE7*-[A]_N_ also exhibited high posterior inclusion probabilities (Table S5) indicating that their effect on hair colour is independent of these well-known biallelic variants. The TRs in this region were weakly correlated with SNP variation in MC1R (Fig. 2B), reinforcing their independent influence on hair colour. When conditioned on one another, the effect of each TR on hair colour persists (Fig. 2C).

**Fig. 2.**
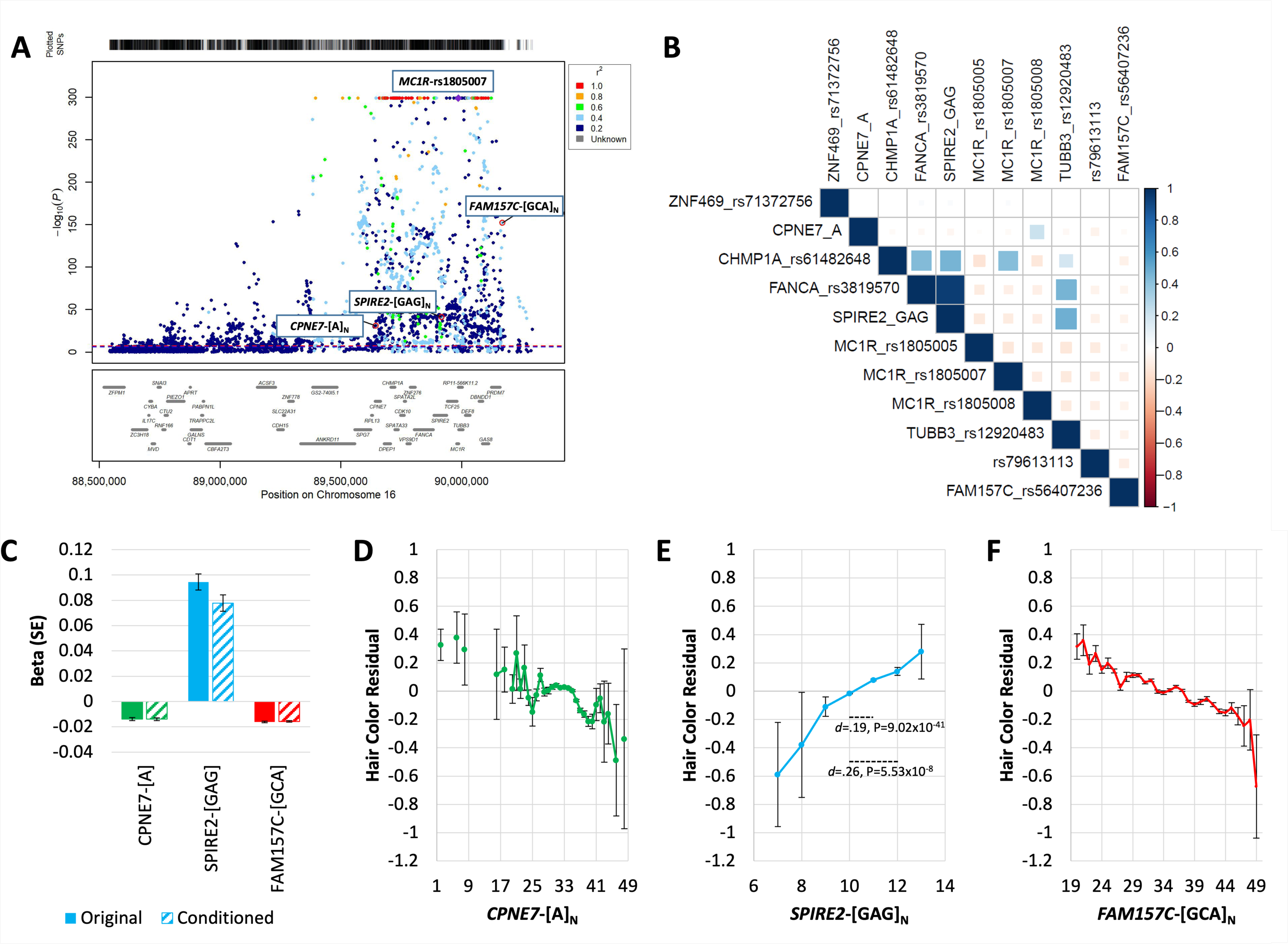
TR effects in the *MC1R* region. (A) Regional plot of the *MC1R*-containing genomic region of chromosome 16. Data points reflect the –log_10_(p-value) for association between each variant and low eumelanin hair colour with the three tandem repeat (TR) signals labeled (all fine-mapping results for each hair colour model are shown in Table S5). For reference to a known pigmentation locus, the *MC1R*-rs1805007 locus also is labeled. In (B), the linkage disequilibrium pattern between the discovered TRs and known pigmentation predictive SNPs is shown. Empty cells in (B) indicate lack of LD (P>0.05) between a pair of variants. Panel C compares the effect of each TR before and after conditioning their effects on the other TRs in the region. In D-F, the relationship between expansion and contraction of the discovered TRs is shown. The likely causal exonic repeat at *SPIRE2* is labeled with measures of effect size compared to the modal genotype using the Cohen’s *d* estimate.

The effects of TR locus-level burden on hair colour are shown in Figures 2D-F. Between the smallest and largest *SPIRE2*-[GAG]_N_ locus-level repeat burden, we observed a large effect on hair colour (Cohen’s *d*=1.16, P=0.031). Relative to the *SPIRE2*-[GAG]_N_ modal genotype (GAG_5_|GAG_5_), carrying an expanded form of the repeat corresponded to a small increase in hair darkening (Cohen’s *d*=0.19, P=9.02×10^-41^ for one additional repeat and *d*=0.26, P=5.53×10^-8^ for two additional repeats; Fig. 2E). Though *SPIRE2*-[GAG]_N_ is exonic and it exhibits a relatively large effect on hair colour across all three models, the mechanism of effect does not relate to gene expression (Table S6), splicing, or 3-dimensional protein folding (Fig. S3).

### Hair colour TRs in the NCOA6 region

A second cluster of significantly associated TRs localized to a region with high gene density on chromosome 20 that contains *NCOA6*, another known pigmentation gene (23). We identified five TRs in this region: *ITCH*-[T]_N_, *PIGU*-[AG]_N_, *TP53INP*-[GT]_N_, *ACSS2*-[GT]_N_, and *RBM12*-[GGCCGG]_N_. Though none of the TRs in this region fine-map with respect to hair colour (Table S5), *TP53INP*-[GT]_N_ associates with splicing quantity in the tibial nerve (slope=0.065, s.e.=0.023, P=0.005, Table S7). Additionally, *RBM12*-[GGCCGG]_N_ encodes a stretch of repeating glycine-proline amino acid residues within a highly disordered region of the resulting protein. We next modeled how the largest and smallest repeat expansions at this position influence protein folding. Expansion and contraction at *RBM12*-[GGCCGG]_N_ had no influence on the local alignment error of the repeat region (Fig. S4). However, the contracted form of *RBM12*-[GGCCGG]_N_ resulted in a large reduction in protein structural confidence (Cohen’s *d*=1.84, P=3.63×10^-5^) within a disordered region of the protein ranging from amino acids 204-210 (mean pLDDT_canonical_=57.21, s.d.=4.68, pLDDT_contracted_=25.15, s.d.=6.18, Fig. S4). The effect in this region was not observed in the expanded form of *RBM12*-[GGCCGG]_N_ (mean pLDDT_expanded_=25.15, s.d.=6.18, P=0.271). Conversely, the expanded form of *RBM12*-[GGCCGG]_N_ resulted in a large reduction in protein structural confidence (Cohen’s *d*=1.78, P=6.26×10^-4^, Fig. S4) within a different disordered region ranging from amino acids 736-740 (mean pLDDT_canonical_=54.43, s.d.=7.13, pLDDT_expanded_=28.86, s.d.=2.20). The effect in this region was not observed in the contracted form of *RBM12*-[GGCCGG]_N_ (mean pLDDT_expanded_=38.84, s.d.=4.28, P=0.061).

### Novel TR-containing regions associated with hair colour

In addition to the detection of loci containing *MC1R* and *NCOA6*, we identified relevant hair colour effects at *SLC45A2*-[AAC]_N_, *FZD*-[CGC]_N_, and *FOXE1*-[GCC]_N_. Though SNP variation within *SLC45A2* is capable of predicting hair colour variation (24), *SLC45A2*-[AAC]_N_ was not part of the credible set of putative causal variants in this locus despite having a relatively large effect on hair colour (blonde hair lJ=0.064, P=1.99×10^-9^, red hair lJ=0.045, P=1.51×10^-9^, low eumelanin lJ=0.051, P=2.49×10^-6^). Relative to the *SLC45A2*-[AAC]_N_ modal genotype (AAC_4_|AAC_4_), the largest effect on low eumelanin hair colour was observed with the addition of a single repeat (Cohen’s *d*=0.15, P=7.95×10^-10^; Fig. S5).

*FOXE1*-[GCC]_N_ encodes an alpha-helical structure composed of alanine residues that, when contracted, results in reduced protein structural confidence (Cohen’s *d*=1.75, P=8.08×10^-4^) within a disordered region of the protein ranging from amino acids 204-210 (mean pLDDT_canonical_=57.21, s.d.=4.68, pLDDT_contracted_=25.15, s.d.=6.18, Fig. 3). This contraction had a significant but relatively small effect in each tested model (blonde hair lJ=0.009, P=1.35×10^-7^, red hair lJ=0.004, P=3.92×10^-5^, low eumelanin lJ=0.009, P=4.63×10^-7^). Though potentially modifying protein 3-dimensional properties, *FOXE1*-[GCC]_N_ did not belong to the fine-mapping credible set for this region of the genome.

**Fig. 3.**
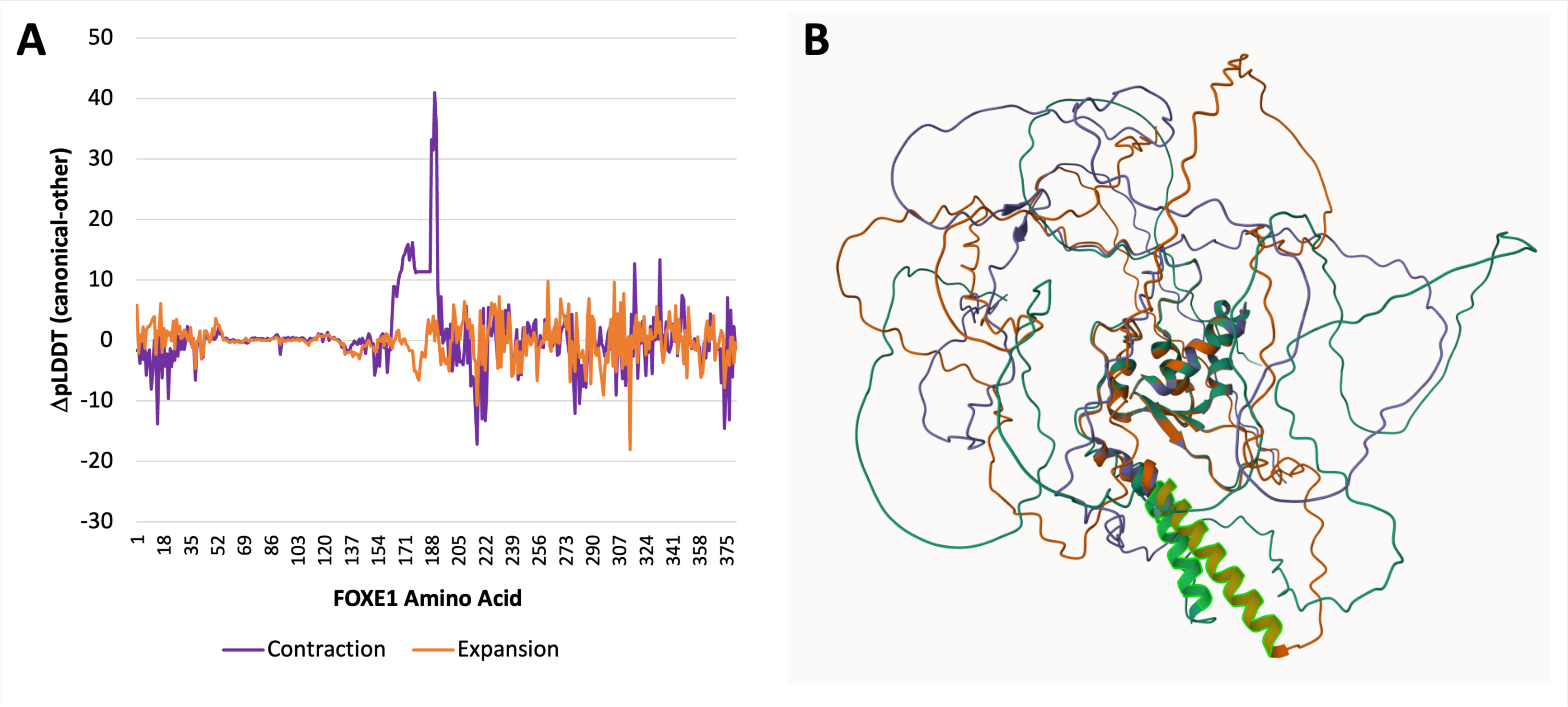
Putative effects of TR variation on FOXE1 structure. (A) Putative impact of tandem repeat mutation on the structure of FOXE1. Change in per-residue folding confidence in canonical FOXE1 was compared to the contraction (in purple) and expansion (in orange). (B) Mol* Viewer was used to superpose the predicted structures with the FOXE1 GCC-containing stretch of alanines highlighted. The purple structure is contracted at *FOXE1*-[GCC]_N_ while the orange structure is expanded at *FOXE1*-[GCC]_N_.

### Additive TR scores associate with darker hair

We next investigated whether the additive effect of TR variation associates with hair colour in 10% of UKB EUR participants excluded from the discovery phase. The 11 TRs discovered from the EUR low eumelanin model localize to 6 independent regions of the genome. We used *CUL2*-[A]_N_, *SPIRE2*-[GAG]_N_, *HGS*-[CCAGCC]_N_, *TP53INP2*-[GT]_N_, *SLC45A2*-[AAC]_N_, and *FOXE1*-[GCC]_N_ to derive polygenic scores (PGS) for hair colour. Effect sizes estimated from the EUR discovery sample were aggregated in a weighted additive model (Equation 1) based on locus level burden of TR repeat size. Our TR PGS associated with hair colour (adjusted R^2^=0.45%, P=1.97×10^-16^) such that relative to the lowest decile of TR PGS, every subsequent decile had significantly higher odds of darker hair. The largest effect was observed between the lowest and highest decile of TR PGS: lJ=0.361, P=9.10×10^-21^ (Fig. 4A).

**Fig. 4.**
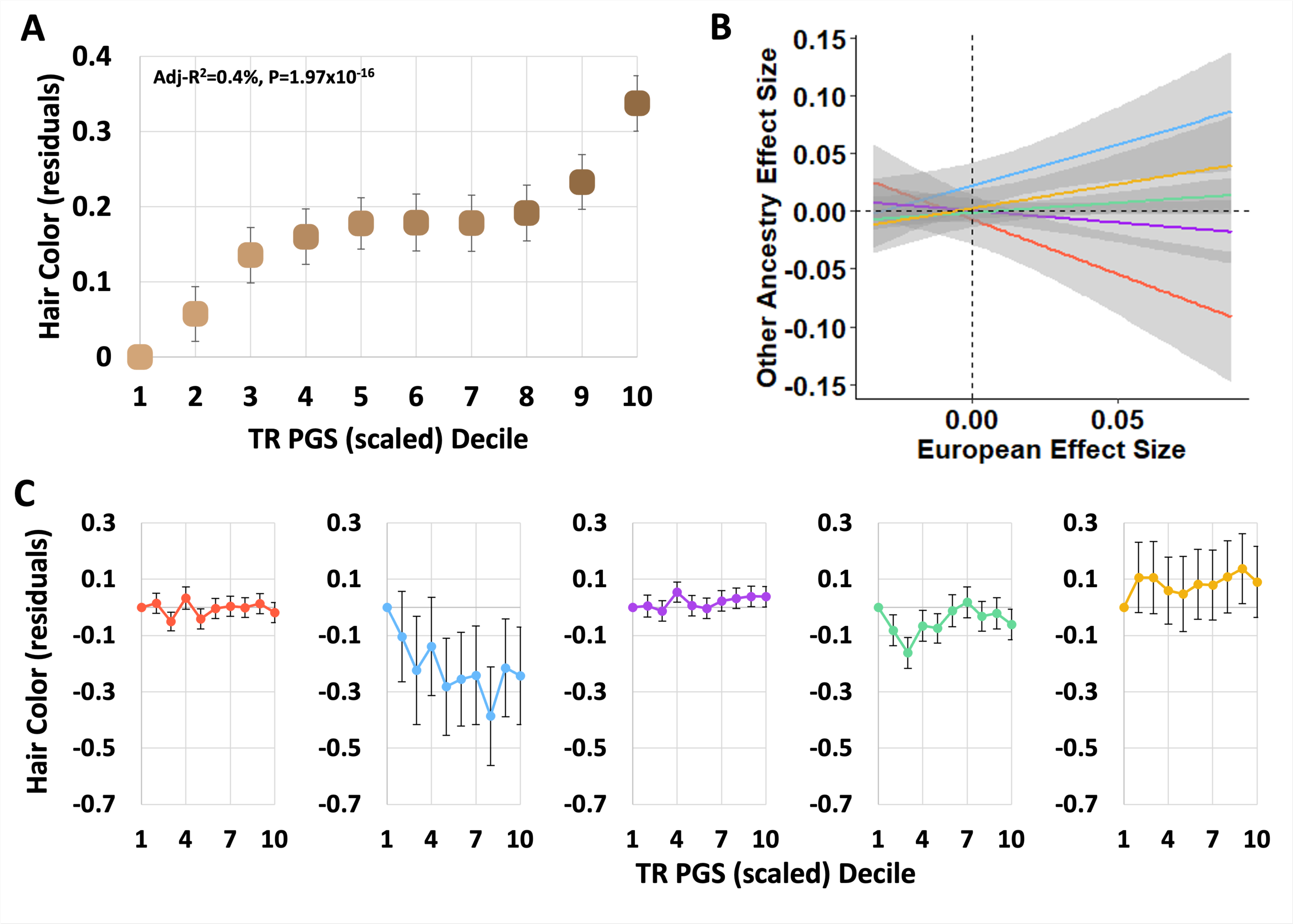
TR polygenic scores (PGS) associate with low eumelanin hair colour. (A) An additive PGS of TR variation using low eumelanin associated TRs associated with hair colour in a subset of European ancestry participants. This PGS distinguishes between light brown and dark brown hair as indicated by the colour of the data points. In Panel (B) European ancestry hair colour effect sizes were associated with effect sizes of the same variants in five other ancestry groups: African in red, admixed American in blue, Central/South Asian in purple, East Asian in green, and Middle Eastern in yellow. In (C), the TR PGS developed in EUR participants was applied to five other ancestry groups.

### TRs and hair colour variation in diverse ancestry groups

Due to the low prevalence of blonde and red hair in diverse ancestry groups, we evaluated TR effects on hair colour in AFR, AMR, CSA, EAS, and MID using the low eumelanin phenotype. Two TRs associated with low eumelanin hair colour in the EUR sample nominally replicated in other ancestry groups (Table S8): *SPIRE2*-[GAG]_N_ in AFR had an opposite effect relative to EUR (lJ=-0.148, P=0.017, P_diff_=0.005) and *FAM157C*-[GCA]_N_ in EAS also had an opposite effect relative to EUR (lJ=0.006, P=0.045, P_diff_=0.005). This minimal cross-population replication was also reflected in the poor translation of the hair colour TR PGS to these groups with adjusted R^2^ values ranging from -0.002 (P=0.691 in AMR) to 0.001 (P=0.143 in EAS; Figs. 4B and 4C).

From exome-wide investigation of low eumelanin hair colour in diverse ancestry groups, we uncovered several novel loci (Table S9). In AFR, we detected *ESAM*-[A]_N_ (lJ=0.0314, P=1.03×10^-5^) but this locus was not part of a fine-mapping credible set indicating. In CSA, *UTP18*-[CGG]_N_ associated with hair colour (lJ=0.217, P=4.26×10^-7^). This variant lies just upstream of UTP18 and is considered a putatively causal variant for hair colour variation based on fine-mapping the region (PIP=0.948). These TRs were not detected in the EUR ancestry analysis (*ESAM*-[A]_N_ lJ=0.002, P=0.285 ; *UTP18*-[CGG]_N_ lJ=-0.016, P=0.173).

## Discussion

Hair colour appears to have a polygenic architecture where different types of genetic variation contribute to its heritability. While GWAS have identified many loci contributing to hair colour genetic architecture (3), common SNPs do not completely explain hair colour variability and meaningful biological consequences are difficult to conclude. We investigated TRs as an important source of genetic variation influencing hair colour and identified 16 TR loci implicated in hair colour variation.

The melanocortin 1 receptor (*MC1R*) gene is well known to be associated with pigmentation of skin and hair, as well as susceptibility to associated cancers (e.g., melanoma) (12). Previous research has identified significant associations between *MC1R* variants and red hair in the UKB cohort (25). These associations are likely in part due to combined effects with other loci, as studying *MC1R* alone has suggested variable penetrance for the red hair phenotype, and accounts for only 73% of hair colour variation in isolation (3). Based on our results, at least two TRs in this region also have independent associations with hair colour in EUR participants. The weak association between our discovered hair colour TRs and known hair colour SNPs supports future work investigating TR variation implicated in other complex traits.

We uncovered a TR in forkhead box E1 (*FOXE1*) that associated with blonde, red, and low eumelanin hair colour with no difference in effect size across hair colour models in EUR participants. This gene encodes thyroid transcription factor 2 (TTF2) which contributes to thyroid function, and in particular, loss-of-function (LOF) mutations have been reported in patients with Bamforth-Lazarus syndrome (26,27). Bamforth-Lazarus syndrome is a congenital condition affecting thyroid function (e.g., hypothyroidism) and has a characteristic symptom of spiky hair observable from birth (26). Previous research has implicated *FOXE1* as necessary for hair follicle morphogenesis via sonic hedgehog signaling, and has noted its LOF as responsible for the spiky hair phenotype in Bamforth-Lazarus syndrome patients (28). Further, one study in mice has suggested inhibition of sonic hedgehog signaling to induce pigmentation via increased expression of melanin (29). Our study noted the *FOXE1*-[GCC]_N_ exonic TR to have high fine-mapping probability and a significant effect on protein folding. This trinucleotide repeat lies just downstream of the *FOXE1* DNA binding domain and has a modest effect on the structural stability of this region, potentially influencing DNA binding efficiency.

This work also revealed novel TRs in other ancestry groups. A TR localized to endothelial cell adhesion molecule (*ESAM*-[A]_N_), involved in the integrin pathway, was significantly associated with hair colour in African ancestry participants (30). Integrin expression influences skin and hair follicle integrity, with induced mutations in this pathway resulting in hair loss in mice (31). Further, previous studies suggested a role for integrins in melanocyte attachment and morphology and therefore may affect pigmentation (32,33). We also detected a TR located in the U3 small nucleolar RNA-associated protein 18 (*UTP18*-[CGG]_N_), a gene implicated in the development of colorectal carcinoma, which was identified in this study in CSA (34). *UTP18* is involved in Hippo signaling, a pathway involved in cell proliferation and for which dysregulation contributes to many disease phenotypes (35). Notably, Hippo pathway failure has been associated with melanoma, and one study has hypothesized this is due to its regulation of melanocyte density in the epidermis (36).

PGS are generally estimated using SNP association statistics for a given trait to assess risk in an independent sample. Recently, however, several studies have demonstrated the feasibility of using structure variants, such as copy number variants (CNVs), to assess additive risk scores for a given trait. For example, a 2022 study generated risk scores using CNVs, which are similar in structure to TRs but encompass a larger repeat segment (37). The study found PGS generated through CNVs to be analogous to SNP scoring methods, and when used in conjunction with SNP polygenic scores improved statistical modeling. We extended this additive scoring logic to investigate the predictive power of an additive TR PGS. Similar to PGS based on SNPs and CNVs, our TR PGS reliably predicted darker hair in the independent EUR sample but had limited predictive power in diverse populations. This observation may be attributed to stark differences in phenotype variation across global population groups and/or poor transferability of hair colour genetics across ancestries. These findings signify the feasibility of using TR-based polygenic scoring to take advantage of different types of genetic architecture that independently account for different aspects of phenotypic variance. We posit that additionally considering SNP loci can further assist in accounting for genetic differences affecting hair colour and melanin production.

This study has several limitations to consider. First, the hair colour phenotype from UKB is a self-report assessment. We therefore rely upon participants to accurately report their hair colour, which is subjective and may be unreliable at intermediate hair colours. A previous study assessing accuracy of self-reported pigmentation phenotypes found only moderate agreement between reported hair colour and spectrophotometer-derived colour (38). To address issues with using discrete measures of pigmentation, recent studies have begun to generate continuous measurements (39,40). One trade-off with using continuous measurements of hair colour may be the misrepresentation of the transition from blonde to brown (capturing eumelanin concentration) *versus* red to brown (capturing both eumelanin and phaeomelanin concentrations). Second, we excluded many variants from our models by focusing on the exome. Though intentional in order to focus on coding variation, the predictive capabilities of intergenic variants may greatly improve model performance in EUR and its transferability to other diverse populations. For example, many non-coding variants located in and around the oculocutaneous albinism II (*OCA2*) gene contribute to hair pigmentation variation, and cannot be identified in the sole study of coding regions (41). Finally, TR variant calling is challenging, with the dozens of available methods yielding unique benefits and drawbacks (42–44). For example, the GangSTR method used herein has an elevated error rate in the context of AC/TG dinucleotide motifs that is not present in other methods that have different shortcomings. Future work will harmonize TR calls from multiple methods for improved genotyping confidence.

In conclusion, we detected and characterized hair colour TRs in EUR, assessing their effect on self-reported hair colour in other ancestry groups. Several of these loci were prioritized relative to surrounding SNP variant and collectively refined our understanding of hair colour and pigmentation biology, most notably through TR variation found in the *MC1R* region. This work adds to an emerging body of literature in support of TR variation impacting common complex traits across various domains.

## Materials and Methods

### Cohort and Hair Colour Description

The UK Biobank is a population-based cohort of >500,000 participants. Participants consented and assessed for behavior, physical appearance, physical activity, medical history, mental health, etc. UKB participants were assigned to one of six population groups using genetic principal components and a random forest classifier based on the 1000 Genomes Project Phase 3 and Human Genome Diversity Panel population groups (45,46). See the Pan-UKB webpage for additional details: https://pan.ukbb.broadinstitute.org on this procedure. Hair colour in the UK Biobank is assessed by Field ID 1747 (“what best describes your natural hair colour (if your hair colour is grey, the colour before you went grey)”). Participants responded with either: “blonde” = “1”, “red” = “2”, “light brown” = “3”, “dark brown” = “4”, “black” = “5”, “other” = “6”. We tested three ordinal models of hair colour. First, we compared blonde, light brown, dark brown, and black with red hair excluded from the model (3). Second, we tested red, light brown, dark brown, and black with blonde hair excluded from the model (3). Third, we combined red and blonde hair into a category of “low eumelanin” hair colour compared to light brown, dark brown, and black hair. All models excluded individuals who reported “other” hair colour. This research has been conducted in the scope of UKB application reference number 58146.

### UK Biobank Tandem Repeat Association Testing

UKB CRAM files of WES reads aligned to the hg38 reference genome were converted to binary alignment map (BAM) files using cramtools v3.0 (February 2021) (47). Genotyping of autosomal tandem repeats from short reads was performed with GangSTR v2.5.0 using sorted and indexed BAMs (48). To associate each multi-allelic TRs with hair colour, TR genotypes were converted to a locus-level burden of allele lengths by summing both TR alleles in a genotype. In R, generalized linear models were used to regress TR burden on hair colour using age, sex, sex×age, age^2^, sex×age^2^, and the first 10 within-ancestry principal components as covariates. The test statistics from TR association models of each trait were evaluated for inflation by estimating lambda in the R package QCEWAS (49). Multiple testing correction was applied using a false discovery rate (FDR) of 5%. We also report the number of loci meeting the genome-wide multiple testing correction common to SNP-based genome-wide association studies (p<5×10^-8^).

Our discovery analysis was performed in 133,190 unrelated European ancestry participants from the UKB. We replicated our European ancestry findings in a subset of 14,799 unrelated participants (10% of the total). In each additional ancestry group, we tested only the model including the “low melanin” hair colour category due to the relatively low prevalence of blonde and red hair colour in these cohorts. GWAS meta-analysis was conducted using the inverse variance weighted method in METAL including AFR (N=2,607), AMR (N=422), CSA (N=3,765), EAS (N=1,153), EUR (N=133,190), and MID (N=640) population groups for a total of up to 141,777 participants (50). Hair colour distributions by population group are shown in Fig. S1.

### Gene set enrichment

Pathway enrichment was performed using Enrichr. Briefly, Enrichr reports three results for a set of genes. The p-value is computed from a hypergeometric test assuming a binomial distribution. The q-value is an adjusted p-value using the Benjamini Hochberg method for multiple testing correction. The odds ratio reflects the representation of input genes in gene sets relative to the proportion of genes that belong to the gene set. These are described in greater detail here: https://maayanlab.cloud/Enrichr/help#background&q=4.

### Gene expression and splicing effects

Gene expression associated TRs (eSTRs) and splicing associated TRs (spl-TRs) were identified using data from Fotsing et al. and Hamanaka et al., respectively (51,52). Both resources used the GTEx Project including all v7 individuals for eSTRs (17 tissues studied in N=652 unrelated donors; 86% European ancestry) and v8 for spl-TRs (49 tissues studied in N=838 unrelated donors; 84.6% European ancestry). For eSTRs, average TR repeat lengths were called from hg19-aligned whole genome sequences using HipSTR (53). For spl-TRs, average TR repeat lengths were called from hg38-aligned whole genome sequences using GangSTR. Both studies used TR dosage as a predictor of expression or splicing quantity. The eSTR models included age, sex, and the first 10 within-ancestry principal components as covariates. The spl-TR models included sex, library preparation protocol, sequencing platform, the first five within-ancestry principal components, and probabilistic estimation of expression residuals.

### Fine-mapping of TR-containing genomic regions

Candidate causal variants were detected using the program FINEMAP (version 1.4.2) (22). FINEMAP employs a Bayesian framework that leverages summary association data and linkage disequilibrium (LD) among variants to calculate the posterior probability of causality using a shotgun stochastic search algorithm. Regions selected for fine-mapping were defined as ± 2Mb on either side of the TR of interest. LD matrices and variant effect sizes were calculated using the same set of UKB individuals used for TR association testing. For each locus, the maximum number of causal variants was set to 20 (i.e., a maximum of 20 credible sets). A credible set is comprised of variants that cumulatively reach a probability of at least 95%. The variants within a credible set are referred to as candidate causal variants and each of them has a corresponding posterior inclusion probability (PIP). FINEMAP results were filtered by removing candidate causal variants with a log_10_ of Bayes Factor (BF) > 2 and p-value < 10^-5^ from each of the 95% credible sets. The Bayes Factor compares the likelihood of two competing hypotheses: 1) the variant is causally associated with the phenotype, and 2) the variant is not associated with the phenotype. Thus, the log_10_BFs represent the strength of evidence supporting one hypothesis over the other; values greater than 2 provide robust evidence of causality.

### 3-dimensional protein structural changes

To better understand the consequences of TR variation, we took advantage of the relationship between protein structure and function. We hypothesized that TRs that modify protein structure likely also influence protein function. Default parameters in AlphaFold v2.2.016 were applied to amino acid string sequences to predict the structure of a subset of proteins containing exonic TRs with fine-mapping probabilities >0.95 (54). We predicted protein structure given the shortest and longest TR allele lengths with frequencies >1% in the European ancestry population from UKB. Per-residue predicted local distance difference test (pLDDT) scores for expanded and contracted proteins were compared to the canonical sequence (accessed via UniProt) to identify changes in local protein folding confidence (55). The pLDDT is a per-residue confidence metric that evaluates local distance differences of all atoms in a model, including stereochemical plausibility. As a measure of local model quality, pLDDT less than 50 is considered a strong predictor of structural disorder (56).

### Polygenic scoring

TRs associated with hair colour were used to construct a PGS based on additive contribution of independent variants. When multiple TRs in the same region were discovered in our analysis, we included (i) the fine-mapped TR or the (ii) largest effect size TR from the region. PGS were calculated as follows:

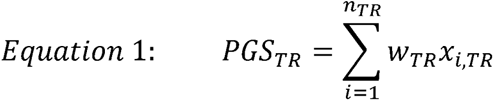

In equation 1, *n_TR_* is the number of TR loci contributing the PGS calculation, *x_i,TR_* is the sum of allele lengths at each TR locus (i.e., the locus-level burden), and *w_TR_*is the effect of the TR on low eumelanin hair colour. To account for linkage disequilibrium, the 11 EUR-significant TRs were reduced to six for PGS generation, removing TRs from the same chromosome. Among TRs on the same chromosome, the TR with the most significant effect size p-value was selected resulting in the following TRs included in the PGS: chr5_33963627, chr9_97854419, chr10_35025203, chr16_90102287, chr17_81700634, and chr20_34709161). TR effect sizes were estimated using a subset of 90% of the unrelated EUR ancestry participants in UKB and tested in the remaining 10% of unrelated EUR participants. R version 4.3.0 and package *Hmisc* were used to generate a 10-decile model for polygenic scoring using ordinary least squares regression. Hair colour residuals per ancestry group (accounting for age, sex, age-by-sex, and the first 10 within-ancestry principal components as covariates) were used in PGS models.

## Acknowledgments

We want to thank all participants from the UK Biobank and the All of Us cohort for their study participation.

## Funding

University of Toronto Data Sciences Institute (DAGY2R1P2 to FRW, NMMN)

University of Toronto Data Sciences Institute (PRSY2R1P05 to FRW)

National Institutes of Health (RF1MH132337 and R33DA047527 to RP)

OneMind (RP)

## Author contributions

Conceptualization: NMMN, FRW

Methodology: MAP, CLA, RP, FRW

Investigation: VF, AM, FRW

Visualization: VF, AM, FRW

Supervision: NMMN, FRW

Writing—original draft: VF, AM, FRW

Writing—review & editing: VF, AM, MAP, CLA, RP, NMMN, FRW

## Data and materials availability

TR-hair colour summary association statistics are available for download at Zenodo: 10.5281/zenodo.10356550. UKB whole exome sequencing data are available to bona fide researchers through approved data access protocol. All other data are available in the main text or the supplementary material.

## Supplementary Materials

**Fig. S1.**
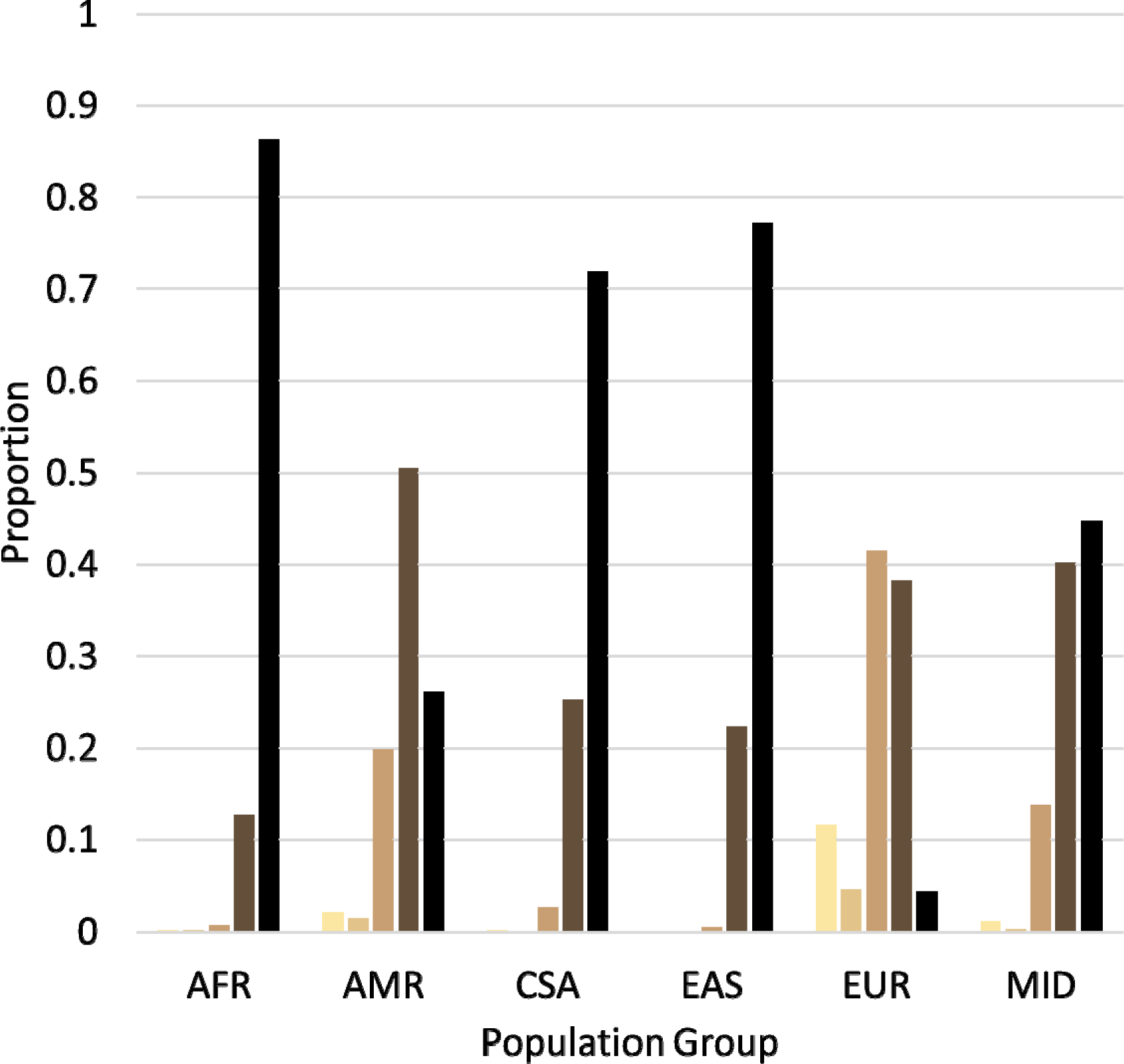
Distribution of hair colour across six ancestry groups in the UK Biobank.

**Fig. S2.**
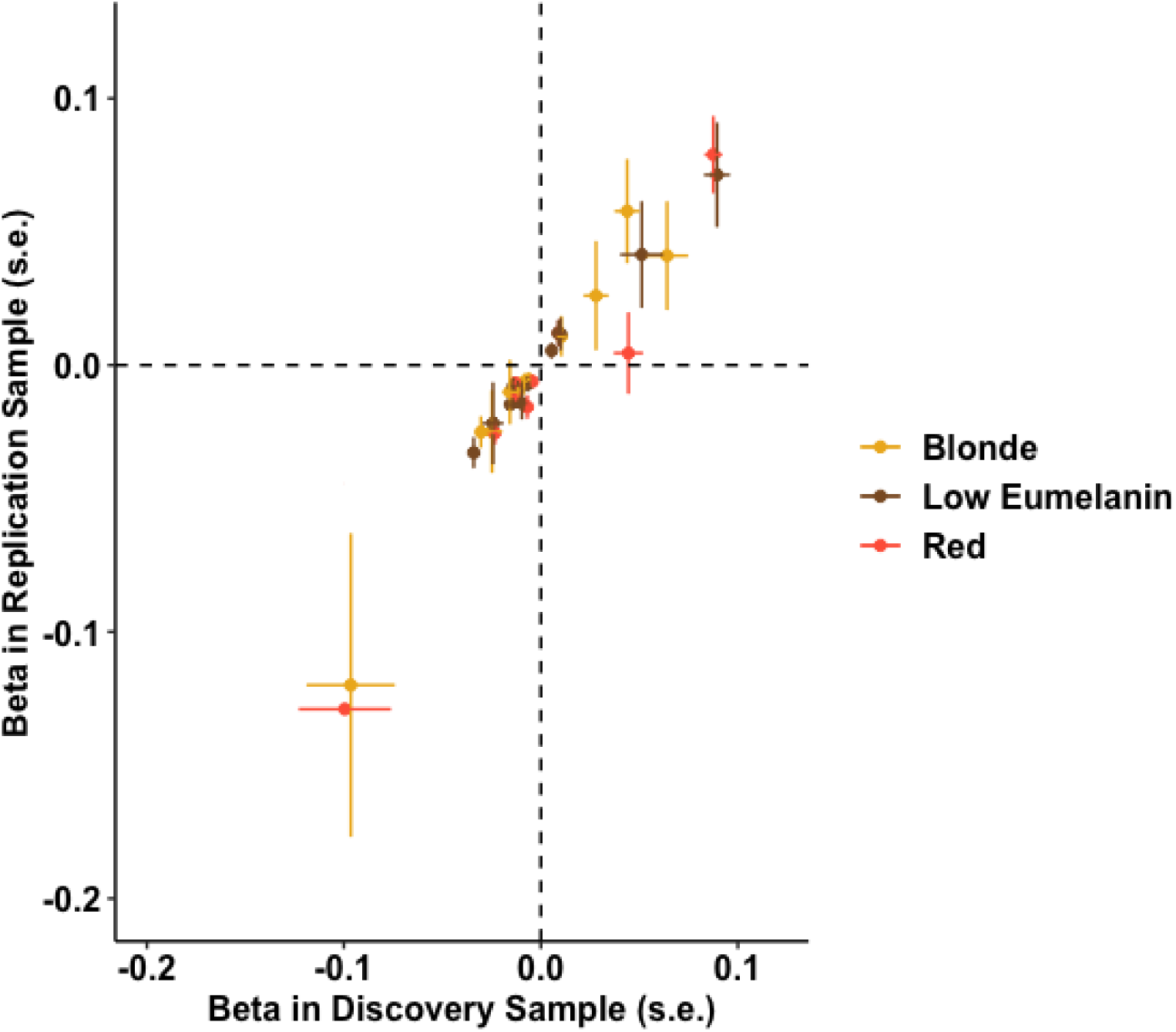
Replication of locus effect sizes in discovery and replication samples of European ancestry participants. Beta estimates and p-values are shown in Table S1.

**Fig. S3.**
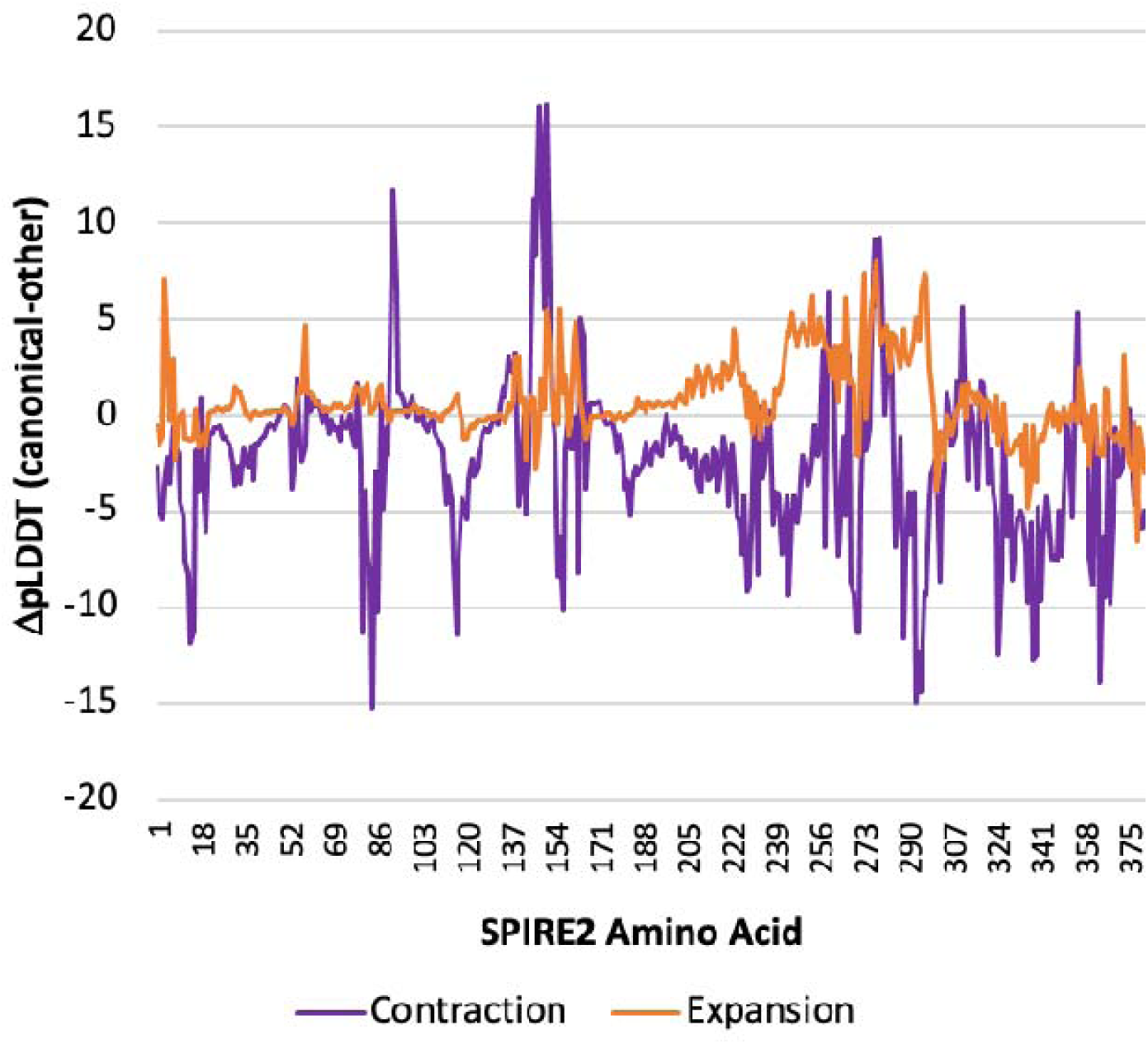
Putative impact of tandem repeat mutation on the structure of SPIRE2. Change in per-residue folding confidence in canonical SPIRE2 was compared to the contraction (in purple) and expansion (in orange).

**Fig. S4.**
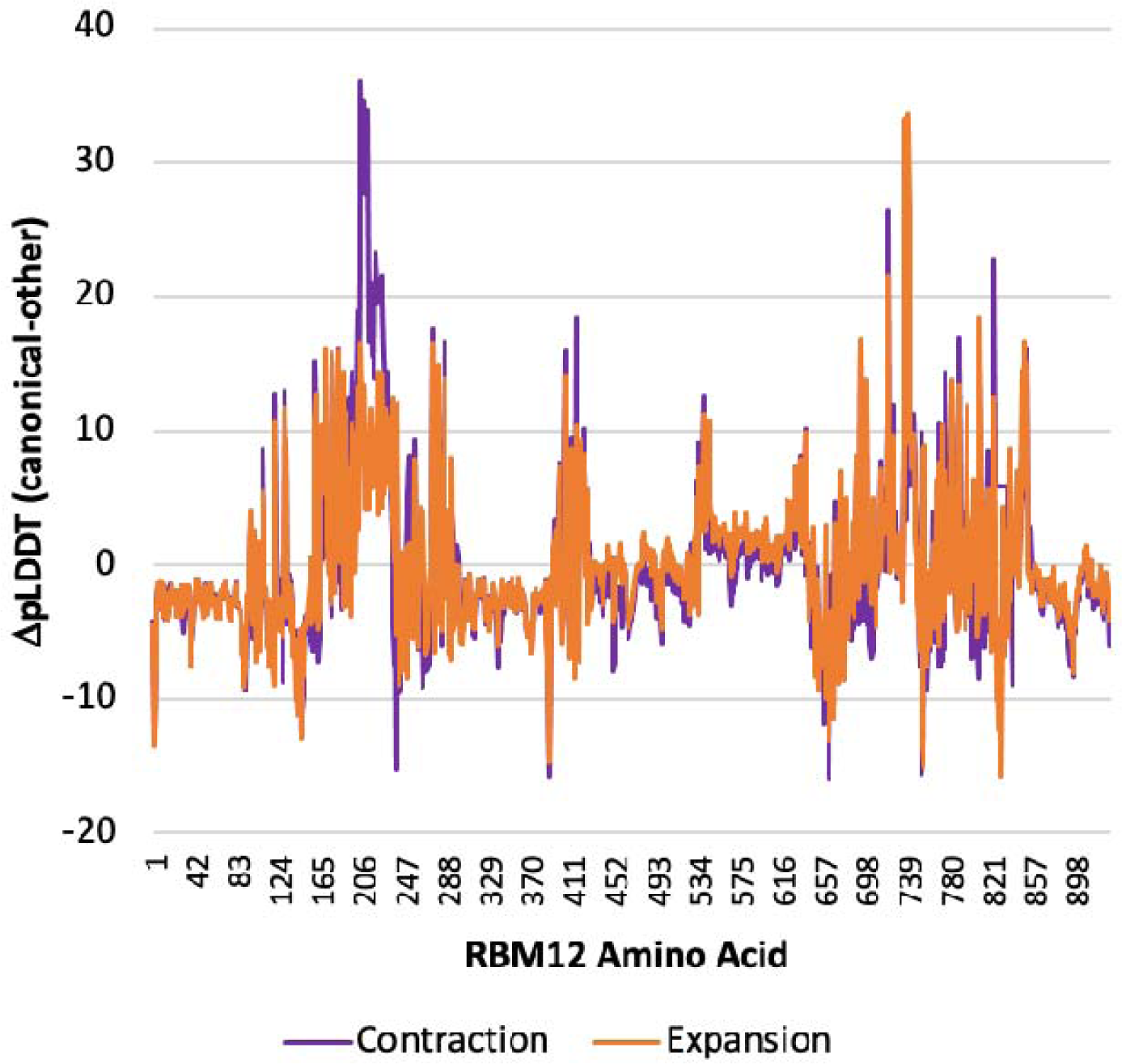
Putative impact of tandem repeat mutation on the structure of RBM12. Change in per-residue folding confidence in canonical RBM12 was compared to the contraction (in purple) and expansion (in orange).

**Fig. S5.**
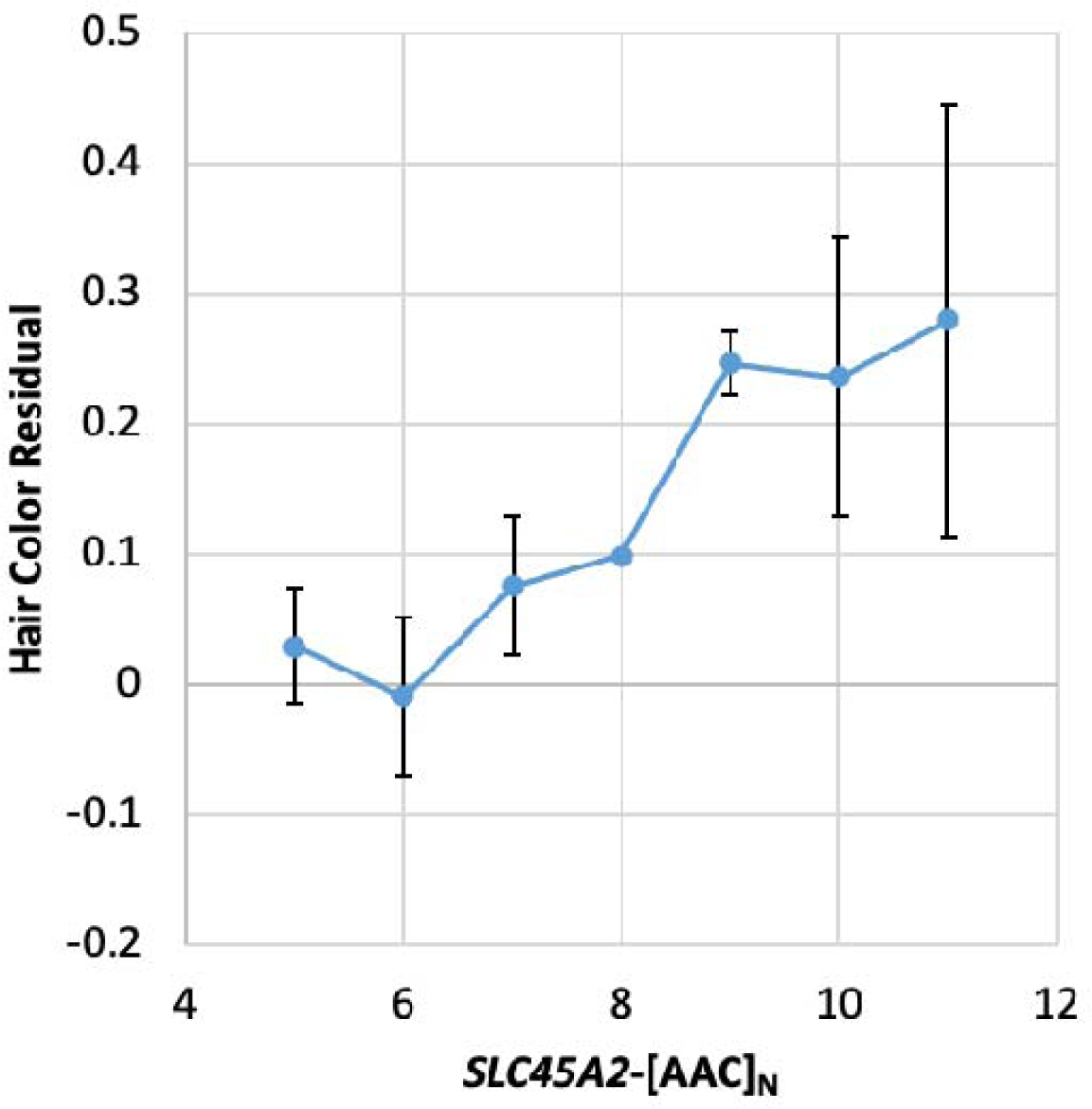
Effects of TR length burden on hair color based on *SLC45A2* genotype. Error bars reflect the standard error.

**Table S1.** Summary statistics for 16 TRs associated with hair colour in European ancestry participants from the UK Biobank. Full summary statistics for 15,947 TRs tested in all three models are available for download at 10.5281/zenodo.10356550.

**Table S2.** Biological process enrichment results from Enrichr for blonde, red, and low eumelanin hair colour models. For each model, the p-value was computed using a Fisher’s exact test and the q-value is an adjusted p-value using the Benjamini-Hochberg method for correction for multiple hypotheses testing. The odds ratio and combined score calculations are described here: https://maayanlab.cloud/Enrichr/help#background&q=4.

**Table S3.** Cellular compartment enrichment results from Enrichr for blonde, red, and low eumelanin hair colour models. For each model, the p-value was computed using a Fisher’s exact test and the q-value is an adjusted p-value using the Benjamini-Hochberg method for correction for multiple hypotheses testing. The odds ratio and combined score calculations are described here: https://maayanlab.cloud/Enrichr/help#background&q=4.

**Table S4.** Molecular function enrichment results from Enrichr for blonde, red, and low eumelanin hair colour models. For each model, the p-value was computed using a Fisher’s exact test and the q-value is an adjusted p-value using the Benjamini-Hochberg method for correction for multiple hypotheses testing. The odds ratio and combined score calculations are described here: https://maayanlab.cloud/Enrichr/help#background&q=4.

**Table S5.** Fine-mapping summary including number of variants per credible set (“N CS”), posterior inclusion probability (PIP), and whether the TR in that region had the highest PIP.

**Table S6.** All expression associated TRs from the SPIRE2 locus. Data retrieved from Fotsing, et al.

**Table S7.** Effect of *TP53INP*-[GT]N on splicing quantity in the tibial nerve. Data retrieved from Hamanaka, et al.

**Table S8.** Cross-population replication of low eumelanin TR effects. Nominally significant replication is highlighted in green.

**Table S9.** Hair colour associated TRs in diverse ancestry groups. Highlighted effects were associated with low eumelanin hair color after multiple testing correction using a false discovery rate of 5% applied per population.

